# Distinct vascular genomic response of proton and gamma radiation

**DOI:** 10.1101/460766

**Authors:** Ricciotti Emanuela, Dimitra Sarantopoulou, Gregory R. Grant, Jenine K. Sanzari, Gabriel S. Krigsfeld, Amber J. Kiliti, Ann R. Kennedy, Tilo Grosser

## Abstract

**Purpose**. The cardiovascular biology of proton radiotherapy is not well understood. We aimed to compare the genomic dose-response to proton and gamma radiation of the mouse aorta to assess whether their vascular effects may diverge.

**Materials and methods.** We performed comparative RNA sequencing of the aorta following (4 hrs) total-body proton and gamma irradiation (0.5 - 200 cGy whole body dose, 10 dose levels) of conscious mice. A trend analysis identified genes that showed a dose response.

**Results.** While fewer genes were dose-responsive to proton than gamma radiation (29 vs. 194 genes; *q*-value ≤ 0.1), the magnitude of the effect was greater. Highly responsive genes were enriched for radiation response pathways (DNA damage, apoptosis, cellular stress and inflammation; *p*-value ≤ 0.01). Gamma, but not proton radiation induced additionally genes in vasculature specific pathways. Genes responsive to both radiation types showed almost perfectly superimposable dose-response relationships.

**Conclusions.** Despite the activation of canonical radiation response pathways by both radiation types, we detected marked differences in the genomic response of the murine aorta. Models of cardiovascular risk based on photon radiation may not accurately predict the risk associated with proton radiation.

## Introduction

Radiotherapy is a widely used cancer treatment resulting in the exposure to ionizing radiation of nearly half a million Americans every year. Therapeutic gamma irradiation that includes the heart and aortic arch in the radiation field is associated with increases in the rates of myocardial infarction, congestive heart failure, valve disease and arrhythmia (1-3). These complications may have long latency times but continue to rise over decades after the initial treatment (4-6) in a radiation dose-dependent fashion (7-9). The dose-response relationship for major cardiac events, such as myocardial infarction, is linear and appears to have a threshold dose to the heart in childhood cancer survivors (10). In breast cancer survivors the rate of major coronary events increased linearly without an apparent threshold dose and was independent of preexisting cardiovascular risk factors (6).

While the underlying molecular mechanisms of gamma radiation-induced cardiovascular disease are not fully understood, inflammatory responses (11, 12) are thought to be an important common feature of the enhanced likelihood of thrombosis (13), accelerated atherosclerosis (11, 14), and impaired cardiac function (15). Gamma radiation-induced experimental atherosclerosis is characterized by vessel wall lesions rich in inflammatory cells (11, 12, 16). Other processes driving the vascular pathology are thought to involve endothelial damage (14, 15), induction of apoptosis (17, 18) and premature cellular senescence (19).

Proton beam therapy has emerged as an alternative to gamma radiotherapy for the treatment of some types of cancer. Its therapeutic use is motivated primarily by an inverted depth-dose profile, the so-called Bragg peak; the proton stops at a specific tissue depth determined by its energy (20). These physical proprieties of proton beams can be exploited to reduce exposure of healthy tissue, such as the heart and the vasculature, by targeting the administered dose more specifically to the tumor (21, 22). While reducing exposure of heart and blood vessels can be reasonably expected to translate into a decrease in acute and chronic toxicity compared with photon radiation (23-26), the dose response relationship between proton irradiation and cardiovascular complication rate has not been established. Prospective investigations comparing the cardiovascular effects of proton beam therapy with conventional photon irradiation have not yet been reported and it is currently unknown whether protons and photons induce similar pathological mechanisms in cardiovascular tissue despite their distinct physics.

Generally, the molecular response following proton radiation-exposure is less well characterized than that of gamma radiation-exposure, because of the limited availability of proton beams for research on model organisms. A small number of studies have directly compared the biological effects of proton and gamma radiation in vivo, but they did not focus on the vasculature. Mice and ferrets respond with a dose-dependent reduction of peripheral blood cell counts to both proton radiation and gamma radiation (27-29). However, gene expression analysis revealed that the molecular changes associated with the apoptotic response varied greatly between proton and gamma radiation in a tissue- and dose-dependent manner (30). Gamma radiation uniquely triggered a stress-response that mediates apoptosis partially independent of the extent of DNA damage. In contrast, proton radiation was associated with increased DNA damage and DNA damage-repair in comparison to exposure to gamma radiation (30). Differences between the radiation types in their effect on gene expression may translate into functional differences. For example, in a three dimensional tissue culture model of endothelial tube formation, protons had a more pronounced dose-dependent effect on vessel structure than gamma photons at equal physical doses (31).

Thus, we hypothesized that the distinct physical interactions of photon and proton radiation with living cells and/or distinct dose response relationships differences might result in detectable differences in the genomic response in blood vessels in vivo. We performed a comparative transcriptome analysis of the early (4 hrs) dose response of the mouse aorta to proton and gamma radiation. While both radiation types activated the core pathways of the early cellular radiation response, we detected marked differences in the genomic response. Thus, it seems plausible that the downstream pathological processes initiated in blood vessels by the induction of gene expression may differ between protons and photons in quality and timing.

## Materials and Methods

**Mice.** Ten to twelve week old male C57B1/6 mice (Jackson Laboratory, ME) were housed in a controlled environment with regard to light, temperature and humidity in the animal facility of the University of Pennsylvania. All mice had free access to food and water. Mice were euthanized through carbon dioxide induced asphyxiation following radiation exposure. The animal care and treatment procedures were approved by the Institutional Animal Care and Use Committees of the University of Pennsylvania.

**Proton and gamma irradiation.** Mice were exposed to ten densely spaced total-body doses of gamma radiation or high energy protons: 0 cGy, 5 cGy, 10 cGy, 25 cGy, 50 cGy, 75 cGy, 100 cGy, 125 cGy, 150 cGy, 200 cGy (N=10 mice per radiation type, one mouse per dose level). This is a more efficient experimental design than using fewer dose levels with multiple replicates per dose level (see ‘Statistical Analysis’). Proton radiation was performed using a proton beam produced by the University of Pennsylvania IBA cyclotron system at a dose rate of 0.5 Gy/min. Total-body gamma radiation was performed from a 137Cs gamma source (Shepherd Mark I Irradiator) at the University of Pennsylvania at a dose rate of 39.25 cGy/min. All mice were irradiated restrained in custom-designed plexiglass chambers. Sham irradiated control mice (0 cGy) were restrained in plexiglass chambers and placed in the gamma or proton irradiators, but they were not irradiated. Proton and photon doses were administered at the same time of the day within a 6 hour time window and in randomized order. There was no difference in the body weight between mice exposed to the gamma or proton radiations (25.4±0.6 *vs* 26.7±0.5 gr, respectively in mice irradiated with gamma or proton radiations). All animals were sacrificed four hours following irradiation. Thoracic aorta, liver, heart and kidney were quickly excised while flushing the thorax and abdominal cavity with ice cold phosphate buffered saline and snap-frozen in liquid nitrogen.

**RNA Sequencing.** Total RNA from aortas were isolated using Trizol and Qiagen RNeasy and the RNA integrity was checked on an Agilent Technologies 2100 Bioanalyzer. RNA-seq of 20 samples on Illumina HiSeq2500 system was performed, using the Illumina TruSeq RNA Sample Preparation Kit and SBS Kit v3. Samples were handled in a blinded fashion during the library preparation and sequencing process. Ribosomal RNA was depleted using a polyA selection protocol.

**RNA-Seq Analysis.** Raw RNA-seq reads were aligned to the mouse genome build mm9 by STAR version 2.5.2a (32). The dataset contained about 6,416,284 sense and 47,258 antisense paired-end stranded 100bp reads, per sample. Data were normalized and quantified at both gene and exon-intron level, using a resampling strategy implemented in the PORT pipeline v0.8.2a-beta (33). A trend analysis (as described in the Statistical Analysis section below) was performed to identify genes that showed a dose response.

**Quantitative Reverse Transcriptase (RT-) PCR.** Total RNA from various tissues (lung, liver, heart and kidney) was isolated using the Trizol and Qiagen RNeasy Kit. Reverse transcription was performed using an RNA-cDNA kit (Applied Biosystems, Carlsbad, CA). Real-time PCR was performed using ABI Taqman primers and reagents on an ABI Prizm 7500 thermocycler according to manufacturer’s instructions. The following primers were used: apoptosis enhancing nuclease (Aen, Mm00471554_m1), cyclin-dependent kinase inhibitor 1a (Cdkn1a, Mm00432448_m1), epoxide hydrolase 1 (Ephx1, Mm00468752_m1) and solute carrier family 19 member 2 (Slc19a2, Mm01290461_m1). All mRNA measurements were normalized to GAPDH mRNA levels (Mm99999915_g1).

### Statistical Analysis

Frequency distribution of the differences between ranks of gene expression were plotted to visualize global differences between radiation types at each dose level. The genes were sorted by descending expression value, ranked by row number, and sorted by the difference in ranks between proton and gamma radiation.

The experimental design used 10 densely spaced dose levels with one mouse per dose. The dose response, measured as the expression trend across doses, was the primary outcome. Such design provides greater statistical power in gene expression profiling than fewer dose levels with more replicates per dose level (34). For example, ten dose levels with one mouse each would provide 80% power to detect a correlation between dose and gene expression of r > 0.95 (Spearman) with an uncorrected p value of ~0.0002. However, here we applied a more robust trend analysis to capture a broader dose-response and conducted a permutation based, non-parametric test for slopes significantly different from horizontal. The trend analysis was performed with two statistics: the number of steps in the same direction (up or down), between consecutive levels of radiation and the slope of the line fitted to the data. Significance was assessed with a permutation distribution obtained by permuting the radiation dose levels thousands of times and for each permutation computing the maximum value of the statistics over all genes. By using the maximum values of the statistics, the tail probabilities of the permutation distribution are automatically corrected for multiple testing. The analysis was performed on sense and antisense signal, for both gene and intron levels. We identified the genes with *q*-value ≤ 0.1. The antisense signal showed no significant findings at this level; thus, only sense signal results are reported. Differences between increasing doses (*q*-value ≤ 0.1) were visualized by plotting the empirical cumulative distribution (eCDF) of the gene expression ratio (expression value at each dose in cGy divided by expression value at 0 cGy) as a non-parametric estimator of the underlying CDF (35).

For more targeted comparison between the two radiation types, we identified the intersection of the genes that were highly responsive to increasing doses (e.g. using a filter of *q*-value ≤ 0.1) in both conditions. Furthermore, we performed a dose response analysis for the 19 genes upregulated by both radiation types. Four of these genes were validated with quantitative RT-PCR, in terms of the mean expression for each radiation dose, the radiation type and the cellular localization.

Enrichment analysis was done using the Ingenuity Knowledge Base (www.ingenuity.com). We ranked genes with a dose-dependent increase in expression by their *q*-value for dose-responsiveness (calculated by trend analysis, see above) and performed pathway enrichment analyses on the top 300 genes in each radiation group. Pathways with a *p*-value ≤ 0.01 (by Ingenuity Pathway Analysis) are reported. Raw data were deposited in Gene Expression Omnibus (NCBI) under accession number GSE105266. (**Supplemental Tables 3** and **4**).

## Results

### Vascular gene expression

*Global comparison between proton and gamma radiation.* We studied the comparative dose-response (0.5 - 200 cGy whole body dose) of aortic gene expression four hours following high energy proton or gamma irradiation. As an initial, qualitative comparison, we plotted the frequency distributions of the differences in gene expression ranks between proton and gamma radiation *(Δ expression rank)* at each dose (**Fig 1**). Narrow distributions of *Δ expression rank* indicate that the impact on gene expression of a physical dose is similar between radiation types.

**Fig.1.**
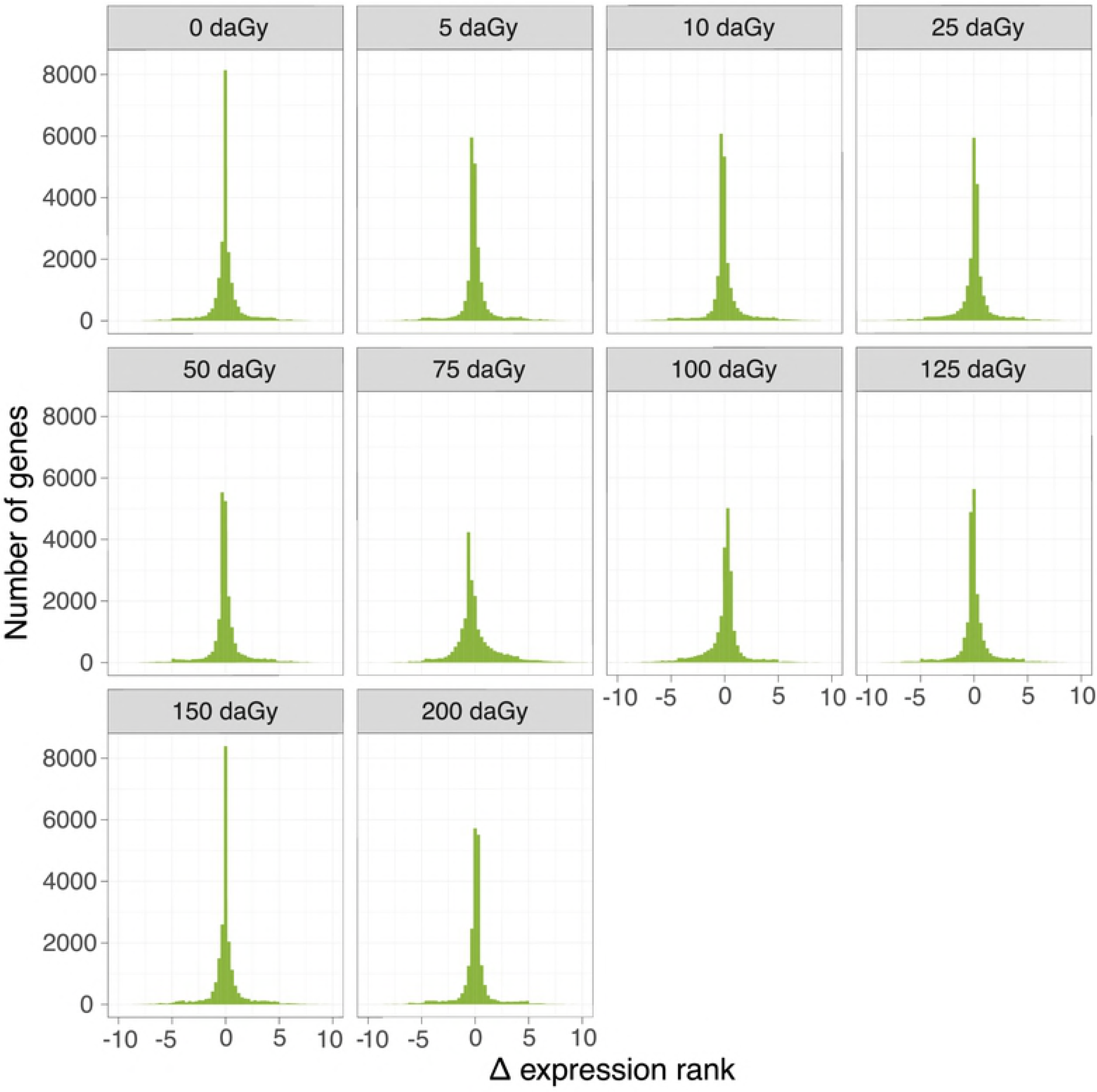
Comparison of the global genomic effects of proton and gamma radiation. Frequency distributions of the difference between expression ranks between proton and gamma radiation were plotted at each physical dose.

*Number of dose responsive genes.* Trend analysis across the 10 doses revealed that fewer genes increased dose-dependently in response to proton radiation than gamma radiation (**Table 1**). At a of *q*-value ≤ 0.1, 29 genes responded with a dose-dependent increase in expression to proton radiation and 194 genes to gamma radiation (**Fig 2** ; **Tables S1 and S2**). A total of 19 genes were upregulated by both types of radiation at this false discovery rate (**Table 2**). We detected no downregulated genes.

**Fig.2.**
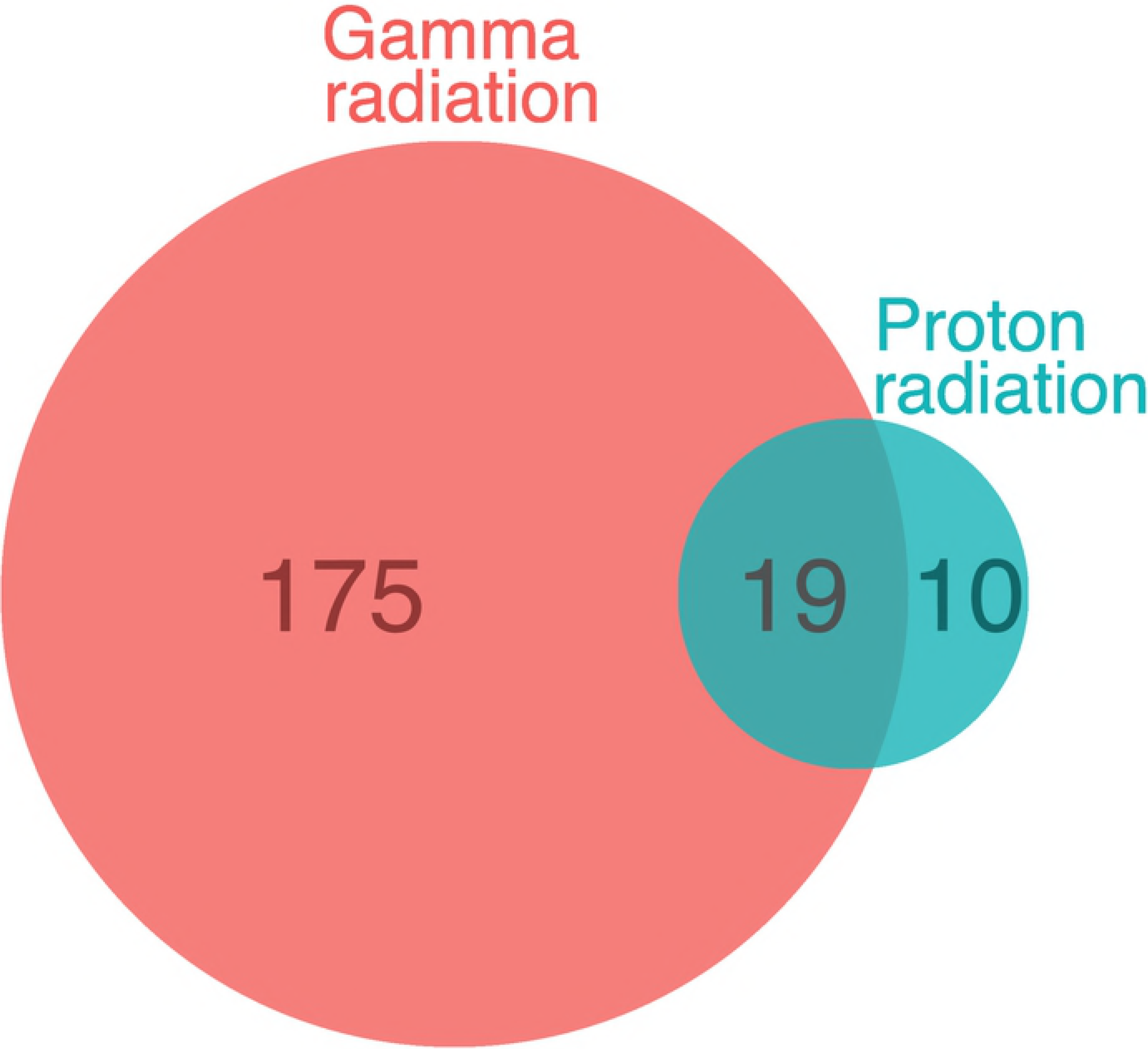
Comparison of the number of dose-responsive genes. A Venn-diagram of genes that present a dose-dependent increase in expression in response to gamma and proton radiation at *q*-value ≤ 0.1 is shown.

**Table 1.** Number of dose-responsive genes at different *q*-value cut-offs.

| $q$-value<br>cut-off | Gamma | | Proton | |
| --- | --- | --- | --- | --- |
|  | # of genes | Average fold change | # of genes | Average fold change |
| **0.5** | 3214 | 1.008 | 218 | 1.011 |
| **0.4** | 2310 | 1.009 | 94 | 1.016 |
| **0.3** | 1578 | 1.009 | 69 | 1.018 |
| **0.2** | 831 | 1.01 | 48 | 1.021 |
| **0.1** | 194 | 1.013 | 29 | 1.024 |
| **0.05** | 54 | 1.014 | 20 | 1.027 |

**Table 2.** Dose-responsive genes to both gamma and proton radiation.

| Gene symbol | Gene name |
| --- | --- |
| Aen | apoptosis enhancing nuclease |
| Dcxr | dicarbonyl L-xylulose reductase |
| Ddx3x | DEAD/H (Asp-Glu-Ala-Asp/His) box polypeptide 3, X-linked |
| Trp53inp1 | transformation related protein 53 inducible nuclear protein 1 |
| Cdkn1a | cyclin-dependent kinase inhibitor 1A (P21) |
| Eif2ak2 | eukaryotic translation initiation factor 2-alpha kinase 2 |
| Ccng1 | cyclin G1 |
| Commd3 | COMM domain containing 3 |
| Ddit4l | DNA-damage-inducible transcript 4-like |
| Gtse1 | G two S phase expressed protein 1 |
| Ephx1 | epoxide hydrolase 1, microsomal |
| Mdm2 | transformed mouse 3T3 cell double minute 2 |
| Cd80 | CD80 antigen |
| Eda2r | ectodysplasin A2 receptor |
| Igf2r | insulin-like growth factor 2 receptor |
| Ano3 | anoctamin 3 |
| Bax | BCL2-associated X protein |
| Slc19a2 | solute carrier family 19 (thiamine transporter), member 2 |
| 9030617O03Rik | RIKEN cDNA 9030617O03 gene |

*Magnitude of the dose-response.* The magnitude of the change in dose-dependent gene expression differed between proton and gamma radiation. Proton radiation caused a more pronounced upregulation on average among the 29 dose-dependent genes than gamma radiation among its 194 dose-dependent genes. This is illustrated by a right shift of the cumulative frequency distribution of proton radiation responsive genes relative to gamma radiation responsive genes (Fig 3). However, a direct comparison of the 19 genes (Table 2) that were responsive to both types of radiation showed that their dose response curves were virtually superimposable (Fig 4). These common genes were amongst those with the most pronounced upregulation. They are involved in various cellular functions (i.e. enzyme, transporter, transmembrane receptor) related to radiation responsive pathways including apoptosis, cell cycle progression and antioxidant defense and distributed across different cellular localization (i.e. nucleus, cytoplasm, plasma membrane).

**Fig.3.**
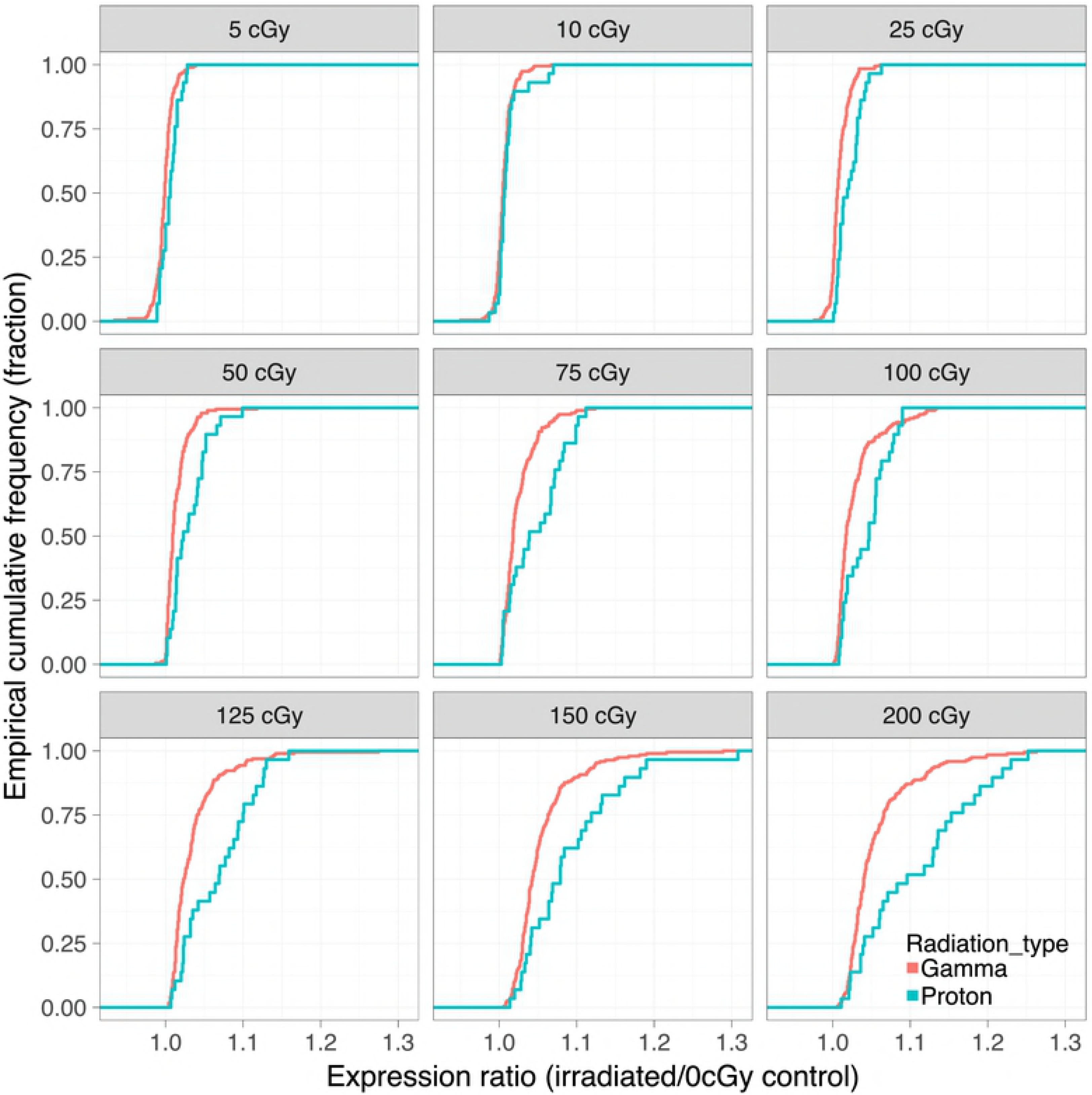
Comparison of the magnitude of the dose response. Empirical cumulative frequency distributions of the gene expression ratios (gene expression at each physical dose over the gene expression at 0 cGy) are plotted for gamma and proton radiation.

**Fig.4.**
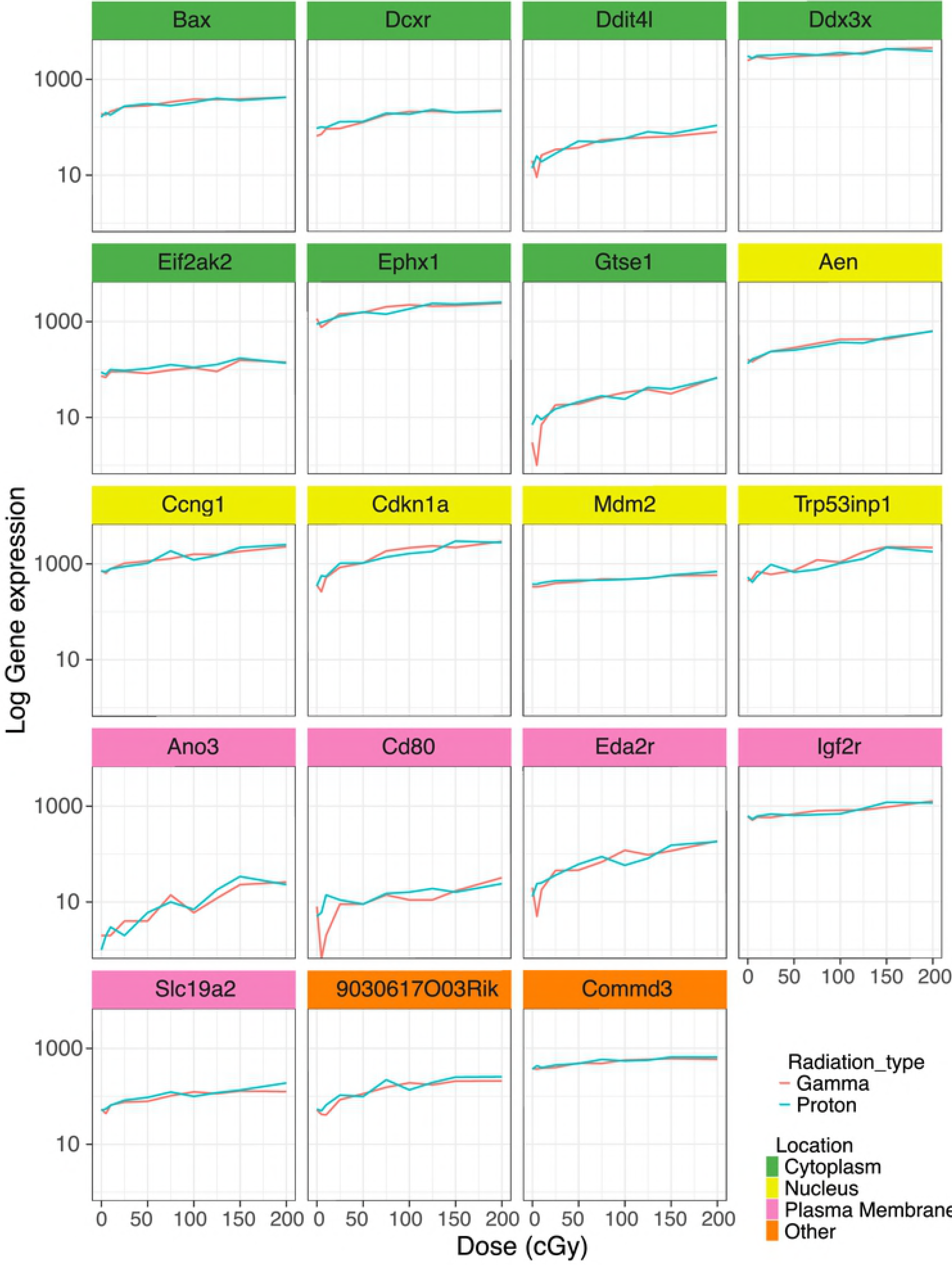
Highly dose-responsive genes upregulated by both proton and gamma radiation. Expression profiles of genes that showed a significant dose-dependent response in the trend analysis (*q*-value ≤ 0.1).

*Validation.* We validated the dose-dependent effects of gamma and proton radiation on the expression of the four most highly responsive genes-Aen, Cdknla, Ephxl and Slc19a2-in the aorta by quantitative RT-PCR. All genes produced a *q*-value of ≤ 0.1 in the trend analysis confirming their dose responsiveness (**Fig 5, top row**). Again, photon and proton induced expression changes were virtually superimposable.

**Fig.5.**
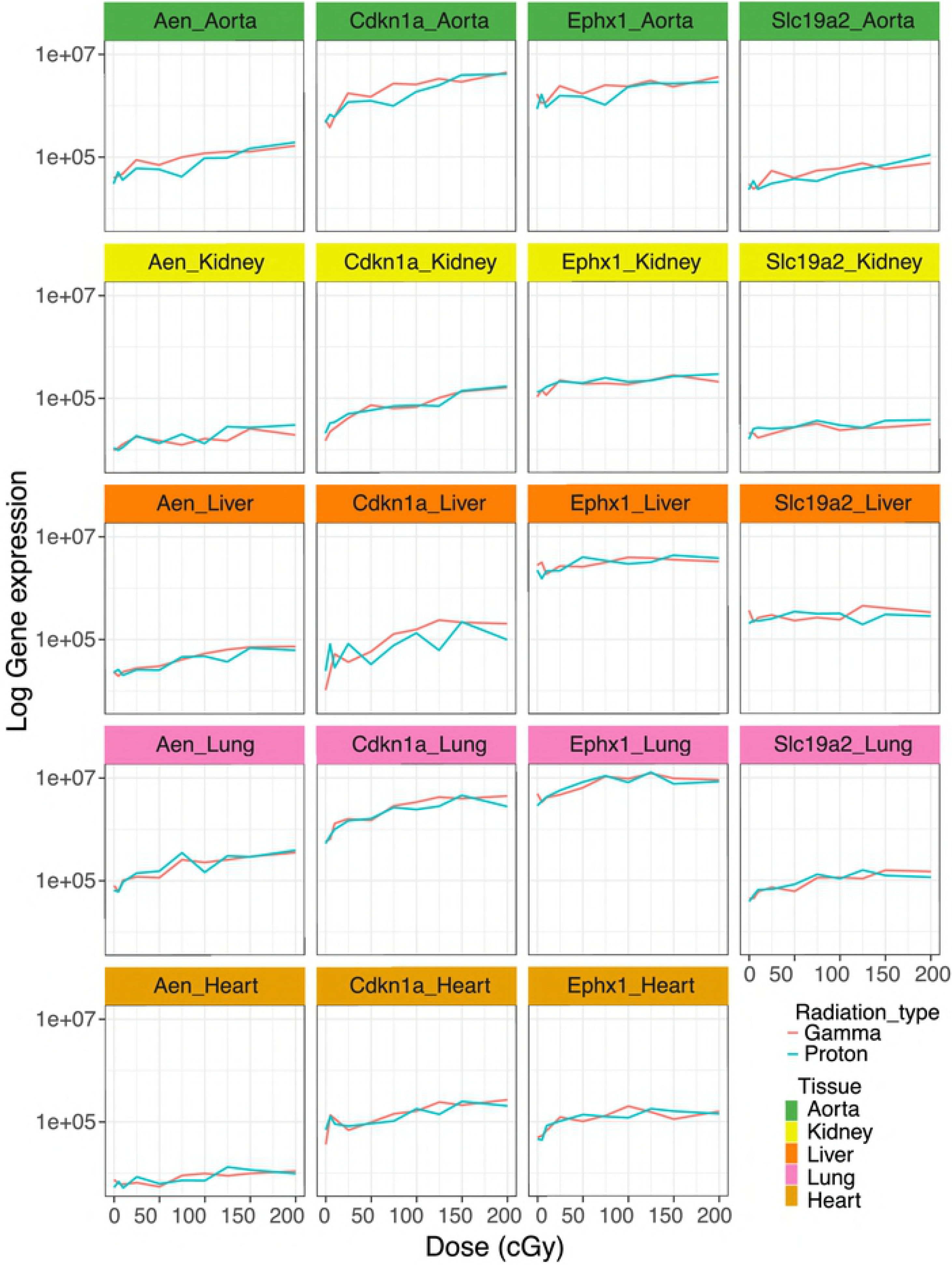
Dose-dependent effects of gamma and proton radiation on Aen, Cdkn1a, Ephx1 and Slc19a2 expression in aorta, kidney, lung and heart. Gene expression was measured by quantitative RT-PCR.

### Dose-response across tissues

We conducted expression analyses on the four most responsive genes-Aen, Cdknla, Ephxl and Slc19a2-also in liver, lung, kidney and heart, to assess whether the similarity in the dose response relationship between proton and gamma radiation was tissue specific. Aen, Cdknla, Ephxl were detectable in these tissues and showed a robust dose-dependent upregulation with a *q*-value of ≤ 0.1 in the trend analysis. Slc19a2 was not expressed in the heart at baseline and we did not observe induction by irradiation (**Fig 5**). Again, the slope of the proton response was virtually identical to that of the gamma radiation response.

### Biological pathways impacted by gamma or proton radiation

The canonical biological pathways enriched for genes that were dose-responsive to gamma or proton radiations (*p* ≤ 0.01) are reported in **Fig 6** . Pathways common to both radiation types were related to p53 dependent apoptosis pathways (p53 signaling, apoptosis signaling, PI3K/AKT signaling, myc mediated apoptosis signaling, aryl hydrocarbon receptor signaling) and p53 independent apoptosis pathways (tumor necrosis factor receptor (TNFR) signaling, granzyme B signaling, signal transducer and activator of transcription 3 (STAT3) pathway, glucocorticoid receptor signaling, death receptor signaling, sumoylation pathway). Both types of radiation also effected DNA damage and cellular stress (cell cycle: G2/M DNA damage checkpoint regulation, ataxia telangiectasia mutated (ATM) signaling and D-glucuronate degradation I) and inflammation (NF-kB and toll like receptor pathways) (**Fig 6**).

**Fig.6.**
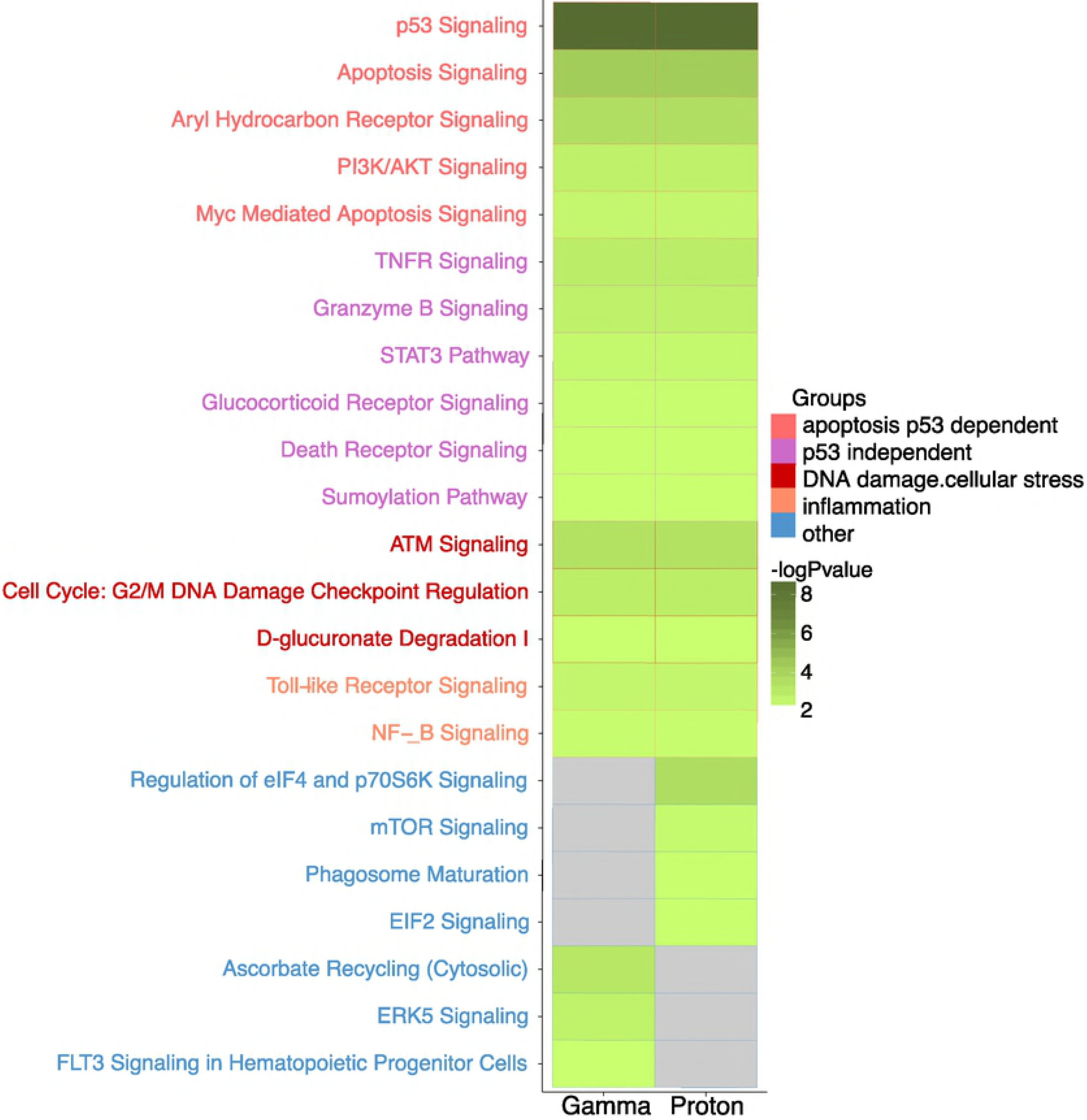
Common canonical pathways enriched by genes that present a dose-dependent increase expression in response to gamma and proton radiation. Although we biased our enrichment analysis against detecting differences by using the 300 most dose-responsive genes of both radiation types regardless of false discovery rate thresholds, we found pathways that were unique to one radiation type. Pathways enriched only by the genes responding to proton radiation were primarily related to cellular growth and stress (eukaryotic initiation factor (eiF) 2 and eiF4 and mechanistic target of rapamycin (mTOR) pathways) and to the cellular immune response (phagosome maturation pathway) (**Fig 6**). Gamma radiation induced a pathway that related to the broader response to oxidative stress (ascorbate recycling pathway) and was not enriched following proton radiation, although individual oxidant stress response genes were clearly upregulated by protons. A vascular process that appeared to be particularly affected by gamma, but not proton radiation was angiogenesis related signaling (extracellular-signal-regulated kinase 5 (ERK5) and Fms like tyrosine kinase 3 (Flt3) pathways) (**Fig 6**).

## Discussion

Radiation induced cardiovascular disease is a recognized sequela of chest photon radiotherapy for conditions such as for mediastinal lymphoma, breast, lung and esophageal cancer (36). The underlying pathophysiological mechanisms involve inflammatory processes in the micro-and macro-vasculature that accelerate atherosclerosis, cause microthrombi and occlusion of vessels, reduced vascular density, perfusion defects and focal ischemia (22, 37, 38). Proton radiotherapy delivers a physical dose in a more targeted fashion than photon irradiation, reducing exposure of the surrounding tissues. However, it is largely unclear whether the biology of photon induced cardiovascular pathologies might similarly apply to proton radiation and adequately sized longterm follow-up studies to determine the cardiovascular hazard associated with proton therapy are not yet available. Gene expression profiles of irradiated tissues were previously shown to correlate with radiation dose (Dressman et al. 2007; Lee et al. 2014; Broustas, Xu, Harken, Garty et al. 2017) and to be predictive of acute radiotherapy-induced adverse effects (42). Furthermore, gene expression profiling comparing relative biological effective-weighted doses of gamma and proton radiation revealed differences in the induction of pro-apoptotic p53-dependent and independent target genes in mice (30). The aim of this study was to compare the vascular genomic response signatures to low doses of proton and gamma radiation administered to conscious animals, in order to predict how similar or dissimilar pathological vascular processes induced by both radiation types might be. This may not only be of relevance in radiation cancer therapy, but also for manned deep-space exploration, which will expose humans to particular radiation, including protons, that does not penetrate the Earth’s geomagnetic shield (43).

We used the aorta as an accessible surrogate tissue for the vascular system and focused on the early molecular radiation effects (4 hours following exposure), which precede the development of structural changes such as intimal hyperplasia or atherosclerosis. We selected a dose range of 0.5 to 200 cGy, which induces dose-dependent effects on the white blood cell counts in mice (29). In human proton beam therapy, the heart and the left anterior descending coronary artery (LAD) is often exposed to doses within this range during the therapy of left-sided breast cancer (44). During photon radiotherapy of patients with breast or chest wall cancer, the heart is usually exposed to higher doses-in the range of 3-17 Gy (total doses given in fractions of 1.8-2.0 Gy)-and the LAD to even higher doses (45). However, epidemiological studies show an increased risk of cardiovascular disease already at markedly lower doses of photon radiation (46-48).

We made the following observations: First, proton radiation resulted in the activation of fewer dose-responsive genes than gamma radiation. For example, six times fewer genes were dose-responsive to proton radiation, when the false discovery rate was set at a *q*-value ≤ 0.1 (**Fig 2, Table 1**). Second, while fewer genes were upregulated by protons, their response was more pronounced on average (**Fig 3**). Proton radiation induced primarily known, highly radiation responsive genes. Similarly, the biological pathways affected by protons included predominantly canonical radiation response functions such as DNA repair, apoptosis, cell growth and inflammation, while gamma radiation induces not only more genes by number, but also a broader range of functions, including for example angiogenesis signaling (**Fig 6**). Third, protons and gamma photons both induced a common set of highly responsive genes, which showed almost perfectly superimposable dose-response relationships (**Fig 4**). We observed the same superimposable dose response relationship of gamma and proton radiations in a subset of genes not only in the aorta but also in liver, lung, heart and kidney (with the exception of Slc19a2 which was not expressed in the heart) (**Fig 5**).

Thus, we found both similarities and intriguing differences in the genomic response to equal physical doses of proton and gamma radiation. Both radiation types induced dose-dependently similar gene sets enriched in the functional categories p53 dependent apoptosis, p53 independent apoptosis, DNA damage, cellular stress and inflammation. DNA lesions induced by ionizing radiation include modifications of the nucleobases, single-strand and doubles strand breaks. The cell responds with activation of repair mechanisms or apoptosis. Thus, the activation of pathways related to p53 dependent apoptosis is consistent with previous reports showing that activation of p53 results in dramatically increased pre-mitotic apoptosis in tissues with a rapid turnover rate such as the hematopoietic system, the gastrointestinal epithelium and endothelial cells (49, 50). Indeed, both high dose of gamma and proton radiations induced a similar number of DNA repair foci in endothelial matrigel cultures, although proton radiation tends to produce larger repair foci, indicating a more complex DNA damage induced by particle proton radiation (51, 52).

The activation of the TNFR signaling pathway, one of the apoptosis p53 independent pathways, has also been shown highly radiation responsive in many tissues and cells (53). Consistent with the fact that inflammatory processes are involved in the initial events triggering atherosclerotic development after radiation exposure, we observed that inflammation associated pathways (NF-kB and Toll like receptor pathways) are sensitive to proton and gamma radiation exposure in a dosage-dependent manner (54). Furthermore, our data confirmed that activation of the ATM kinase pathway is an early event in cellular responses to both gamma and proton irradiation (55, 56).

The dose-dependent expression changes induced by exposure to both, proton or gamma radiation, suggest that at least some of the molecular damage caused in aortic cells *in vivo,* including DNA damage, is similar. Indeed, a previous comparison of higher equivalent doses of gamma and proton radiations show a similar effect of both radiations on pro-apoptotic p53-target genes in the spleens of treated mice (30). However, as mentioned above, in mouse spleen gamma radiation uniquely triggered a pro-apoptotic expression profile while proton radiation triggered a stress-response that mediates apoptosis partially independent of the extent of DNA damage (30). Here, applying lower energy doses, we did not observe this distinction. Both radiation types caused an increased expression of members of the Granzyme B Signaling pathway and Aryl Hydrocarbon Receptor Signaling pathways in the aorta, markers of a response independent of the extent of DNA damage.

In addition to these functional similarities in the response to proton and photon radiations, we also observed similar energy dose response relationships. Thus, the dose response curves of the 19 genes highly responsive to both radiation types were virtually identical. Several well-known radiation responsive genes are among those regulated by both radiation types. *Aen* has been identified as a nuclease that enhances apoptosis following ionizing irradiation (57) and shows dose dependent responses to photon radiation in human blood cells (Broustas, Xu, Harken, Chowdhury et al. 2017) and skin (59). *Cdkn1a* is an inhibitor of G1/S cyclin-dependent kinases that plays a crucial role in the DNA damage signaling in response to radiation (60). Cdkn1a protein expression has been reported to be upregulated in a dose dependent manner both by photon and proton radiation in human fibroblasts (61). Moreover, Cdkn1a gene and protein expression are induced by both gamma and proton radiations in human lens epithelial cells (62). *Ephx1* plays an important role in the detoxification of electrophiles and oxidative stress (63). *Slc19a2* or thiamin transporter THTR1, together with Slc19a3/THTR2, transports thiamin into the cell (64). Slc19a2/THTR1 has been shown to be up-regulated in breast cancer (65) and its expression seems to have a negative effect on tumor specific radiosensitization (66).

We also detected pathways that were differentially activated by both radiation types. Pathways related to cellular growth and cellular stress (eiF2, eiF4 and mTOR pathways) (56) and to cellular immune response (phagosome maturation pathway) were enriched uniquely by proton radiation (28, 67). Pathways related to cell death (ERK5 and Flt3 pathways) and to oxidative stress (ascorbate recycling pathway) were enriched uniquely by gamma radiation. Indeed, microvascular cell death is thought to be an important component of the ischemic injury that initiates radiation-induced inflammatory processes and leads to tissue fibroses (68, 69). Activation of Flt-3 pathway is thought to provide radioprotection to hematopoietic progenitor cells (70, 71) and reactive oxygen species produced by xanthine oxidase following gamma radiation may contribute to endothelial dysfunction and increased vascular stiffness (72). An effect of gamma radiation on ascorbate recycling pathway has not been previously reported.

Proton radiation has been shown to have no effect on or to inhibit angiogenesis related processes while gamma radiation increases expression of angiogenic factors in isolated cells (31,51,73,74). Here-in the adult vasculature-an impact on angiogenic signaling was primarily seen with gamma radiation.

Our study has limitations. The gene expression profiles were generated from male, adult mice from a single strain in response to low doses of a single radiation exposure after a short period of time (acute response). Since the gene expression analyses were done on the whole aorta which contains several cell types, we cannot determine cell type specific changes in gene expression. Irradiation-induced cardiovascular pathologies are noted long (often years) after irradiation therapies. The early gene expression responses detected in our study may not be directly related to such delayed vascular pathologies but may represent early events that could predispose to cardiovascular side-effects.

In conclusion, our RNA sequencing-based expression analysis profiled the changes in aortic gene expression dose response of gamma and proton radiation exposure. Despite the activation of core pathways of the cellular response by both radiation types, we detected marked differences in the genomic response. It seems plausible that these genomic differences may translate into differences in the biological processes leading to cardiovascular pathologies. Thus, our data justify investment in mechanistic research in model organisms, such as models of atherogenesis or vascular injury, to address the potential differential effects of gamma and proton radiation on cardiovascular outcomes.

### Funding

This work was supported by National Institutes of Health Grants HL117798, UL1TR000003, and in part by the Institute for Translational Medicine and Therapeutics of the Perelman School of Medicine at the University of Pennsylvania.

### Author Contributions

E.R.: Investigation, Conceptualization, Methodology, Writing-Original Draft Preparation

D.S.: Investigation, Data Curation, Data Analysis, Writing-Original Draft Preparation

G.R.G. Formal Analysis, Visualization, Writing-Review & Editing

J.S. Investigation, Methodology, Conceptualization, Resources

G.K. Investigation, Methodology, Conceptualization, Resources

A.K. Conceptualization, Methodology, Resources, Writing-Review & Editing

T.G.: Conceptualization, Supervision, Formal Analysis, Visualization, Funding Acquisition, Writing-Original Draft Preparation

## Supplemental Materials Legends

S1 **Table. List of 194 genes that present a dose-dependent increase expression in response to gamma radiation at *q*-value ≤ 0.1.**

S2 **Table. List of 29 genes that present a dose-dependent increase expression in response to proton radiation at *q*-value ≤ 0.1.**

S3 **Table. List of the top 300 genes that present a dose-dependent increase expression in response to gamma radiation sorted by *q*-value.**

S4 **Table. List of the top 300 genes that present a dose-dependent increase expression in response to proton radiation sorted by *q*-value.**

## References

1 Aleman BMP, Van Den Belt-Dusebout AW, De Bruin ML, Van ’t Veer MB, Baaijens MHA, De Boer JP, et al. Late cardiotoxicity after treatment for Hodgkin lymphoma. Blood. 2007; 109(5): 1878–86.

2 Swerdlow AJ, Higgins CD, Smith P, Cunningham D, Hancock BW, Horwich A, et al. Myocardial infarction mortality risk after treatment for Hodgkin disease: a collaborative British cohort study. J Natl Cancer Inst. 2007; 99(3): 206–14.

3 Adams MJ, Lipsitz SR, Colan SD, Tarbell NJ, Treves ST, Diller L, et al. Cardiovascular status in long-term survivors of Hodgkin’s disease treated with chest radiotherapy. J Clin Oncol. 2004; 22(15): 3139–48.

4 Aleman BM, van den Belt-Dusebout AW, Klokman WJ, Van’t Veer MB, Bartelink H, van Leeuwen FE. Long-term cause-specific mortality of patients treated for Hodgkin’s disease. J ClinOncol. 2003; 21(18): 3431–9.

5 Henson KE, McGale P, Taylor C, Darby SC. Radiation-related mortality from heart disease and lung cancer more than 20 years after radiotherapy for breast cancer. Br J Cancer. 2013; 108(1): 179–82.

6 Darby SC, Ewertz M, McGale P, Bennet AM, Blom-Goldman U, Brønnum D, et al. Risk of ischemic heart disease in women after radiotherapy for breast cancer. N Engl J Med. 2013; 368(11): 987–98.

7 Shimizu Y, Kodama K, Nishi N, Kasagi F, Suyama A, Soda M, et al. Radiation exposure and circulatory disease risk: Hiroshima and Nagasaki atomic bomb survivor data, 1950-2003. BMJ. 2010; 340: b5349.

8 Finch W, Shamsa K, Lee MS. Cardiovascular complications of radiation exposure. Rev Cardiovasc Med. 2014; 15(3): 232–44.

9 Little MP. A review of non-cancer effects, especially circulatory and ocular diseases. Radiat Environ Biophys. 2013 Nov;52(4): 435–49.

10 Tukenova M, Guibout C, Oberlin O, Doyon F, Mousannif A, Haddy N, et al. Role of cancer treatment in long-term overall and cardiovascular mortality after childhood cancer. J Clin Oncol. 2010; 28(8): 1308–15.

11 Stewart FA, Heeneman S, te Poele J, Kruse J, Russell NS, Gijbels M, et al. Ionizing Radiation Accelerates the Development of Atherosclerotic Lesions in ApoE-/-Mice and Predisposes to an Inflammatory Plaque Phenotype Prone to Hemorrhage. Am J Pathol. 2006; 168(2): 649–58.

12 Hoving S, Heeneman S, Gijbels MJJ, te Poele JAM, Russell NS, Daemen MJAP, et al. Single-Dose and Fractionated Irradiation Promote Initiation and Progression of Atherosclerosis and Induce an Inflammatory Plaque Phenotype in ApoE-/-Mice. Int J Radiat Oncol Biol Phys. 2008; 71(3): 848–57.

13 Vodovotz Y, Waksman R, Kim WH, Bhargava B, Chan RC, Leon M. Effects of intracoronary radiation on thrombosis after balloon overstretch injury in the porcine model. Circulation. 1999; 100(25): 2527–33.

14 Gabriels K, Hoving S, Seemann I, Visser NL, Gijbels MJ, Pol JF, et al. Local heart irradiation of ApoE-/-mice induces microvascular and endocardial damage and accelerates coronary atherosclerosis. Radiother Oncol. 2012; 105(3): 358–64.

15 Seemann I, Gabriels K, Visser NL, Hoving S, Te Poele JA, Pol JF, et al. Irradiation induced modest changes in murine cardiac function despite progressive structural damage to the myocardium and microvasculature. Radiother Oncol. 2012; 103(2): 143–50.

16 Schiller NK, Kubo N, Boisvert Wa, Curtiss LK. Effect of gamma-irradiation and bone marrow transplantation on atherosclerosis in LDL receptor-deficient mice. Arterioscler Thromb Vasc Biol. 2001; 21(Ldl): 1674–80.

17 Panganiban RA, Mungunsukh O, Day RM. X-irradiation induces ER stress, apoptosis, and senescence in pulmonary artery endothelial cells. Int J Radiat Biol. 2013; 89(8): 656–67.

18 Chen JJ, Shah JL, Harris JP, Bui TT, Schaberg K, Kong CS, et al. Clinical Outcomes in Elderly Patients Treated for Oral Cavity Squamous Cell Carcinoma. Radiat Oncol Biol. 2017; 98(4): 775–83.

19 Yentrapalli R, Azimzadeh O, Barjaktarovic Z, Sarioglu H, Wojcik a, Harms-Ringdahl M, et al. Quantitative proteomic analysis reveals induction of premature senescence in human umbilical vein endothelial cells exposed to chronic low-dose rate gamma-radiation. Proteomics. 2013; 13(7): 1096–107.

20 Tommasino F, Durante M. Proton radiobiology. Cancers (Basel). 2015 Feb 12;7(1): 353–81.

21 Gagliardi G, Constine LS, Moiseenko V, Correa C, Pierce L, Allen AM, et al. Thorax : Heart RADIATION DOSE-VOLUME EFFECTS IN THE HEART. Int J Radiat Oncol. 2010; 76(3): 77–85.

22 Trott KR, Doerr W, Facoetti A, Hopewell J, Langendijk J, Van Luijk P, et al. Biological mechanisms of normal tissue damage: Importance for the design of NTCP models. Radiother Oncol. 2012; 105(1): 79–85.

23 Efstathiou JA, Paly JJ, Lu HM, Athar BS, Moteabbed M, Niemierko A, et al. Adjuvant radiation therapy for early stage seminoma: Proton versus photon planning comparison and modeling of second cancer risk. Radiother Oncol. 2012; 103(1): 12–7.

24 Roelofs E, Engelsman M, Rasch C, Persoon L, Qamhiyeh S, de Ruysscher D, et al. Results of a Multicentric In Silico Clinical Trial (ROCOCO). J Thorac Oncol. 2012; 7(1): 165–76.

25 van de Water TA, Bijl HP, Schilstra C, Pijls-Johannesma M, Langendijk JA. The Potential Benefit of Radiotherapy with Protons in Head and Neck Cancer with Respect to Normal Tissue Sparing: A Systematic Review of Literature. Oncologist. 2011; 16(3): 366–77.

26 Van De Water TA, Lomax AJ, Bijl HP, De Jong ME, Schilstra C, Hug EB, et al. Potential benefits of scanned intensity-modulated proton therapy versus advanced photon therapy with regard to sparing of the salivary glands in oropharyngeal cancer. Int J Radiat Oncol Biol Phys. 2011; 79(4): 1216–24.

27 Kajioka EH, Andres ML, Li J, Mao XW, Moyers MF, Nelson GA, et al. Acute effects of whole-body proton irradiation on the immune system of the mouse. Radiat Res. 2000; 153(5 Pt 1): 587–94.

28 Sanzari JK, Romero-Weaver AL, James G, Krigsfeld G, Lin L, Diffenderfer ES, et al. Leukocyte Activity Is Altered in a Ground Based Murine Model of Microgravity and Proton Radiation Exposure. PLoS One. 2013; 8(8): e71757.

29 Maks CJ, Wan XS, Ware JH, Romero-Weaver AL, Sanzari JK, Wilson JM, et al. Analysis of white blood cell counts in mice after gamma-or proton-radiation exposure. Radiat Res. 2011; 176(2): 170–6.

30 Finnberg N, Wambi C, Ware JH, Kennedy AR, El-Deiry WS. Gamma-radiation (GR) triggers a unique gene expression profile associated with cell death compared to proton radiation (PR) in mice in vivo. Cancer Biol Ther. 2008; 7(12): 2023–33.

31 Grabham P, Hu B, Sharma P, Geard C. Effects of Ionizing Radiation on ThreeDimensional Human Vessel Models: Differential Effects According to Radiation Quality and Cellular Development. Radiat Res. 2011; 175(1): 21–8.

32 Dobin A, Davis CA, Schlesinger F, Drenkow J, Zaleski C, Jha S, et al. STAR: Ultrafast universal RNA-seq aligner. Bioinformatics. 2013; 29(1): 15–21.

33 Kim EJ, Lahens NF, Ricciotti E, Manduchi E, Sarantopoulou D, Tishkoff S, et al. PORT: Pipeline of RNA-Seq Transformations [Internet] [cited 2018 Mar 28]. Available from: https://github.com/itmat/Normalization

34 Sefer E, Kleyman M, Bar-Joseph Z. Tradeoffs between Dense and Replicate Sampling Strategies for High-Throughput Time Series Experiments. Cell Syst. 2016; 3(1): 35–42.

35 Nayak S, Lahens N, Kim EJ, Ricciotti E, Paschos G, Tiskoff S, et al. Iso-relevance Functions-A Systematic Approach to Ranking Genomic Features by Differential Effect Size. bioRxiv. 2018; 381814.

36 Lee PJ, Mallik R. Cardiovascular effects of radiation therapy: practical approach to radiation therapy-induced heart disease. Cardiol Rev. 2005; 13(2): 80–6.

37 Lancellotti P, Nkomo VT, Badano LP, Bergler-Klein J, Bogaert J, Davin L, et al. Expert consensus for multi-modality imaging evaluation of cardiovascular complications of radiotherapy in adults: a report from the European Association of Cardiovascular Imaging and the American Society of Echocardiography. Eur Hear Journal-Cardiovascular Imaging J Am Soc Echocardiogr Eur Hear J-Cardiovasc Imaging. 2013; 14: 721–40.

38 Stewart FA, Hoving S, Russell NS. Vascular damage as an underlying mechanism of cardiac and cerebral toxicity in irradiated cancer patients. Radiat Res. 2010; 174(6): 865–9.

39 Dressman HK, Muramoto GG, Chao NJ, Meadows S, Marshall D, Ginsburg GS, et al. Gene expression signatures that predict radiation exposure in mice and humans. PLoS Med. 2007; 4(4):e106.

40 Lee KF, Weng JTY, Hsu PWC, Chi YH, Chen CK, Liu IY, et al. Gene expression profiling of biological pathway alterations by radiation exposure. Biomed Res Int. 2014; 2014: 834087.

41 Broustas CG, Xu Y, Harken AD, Garty G, Amundson SA. Comparison of gene expression response to neutron and x-ray irradiation using mouse blood. BMC Genomics. 2017; 18(1): 2.

42 Mayer C, Popanda O, Greve B, Fritz E, Illig T, Eckardt-Schupp F, et al. A radiation-induced gene expression signature as a tool to predict acute radiotherapy-induced adverse side effects. Cancer Lett. 2011; 302(1): 20–8.

43 Hughson RL, Helm A, Durante M. Heart in space: effect of the extraterrestrial environment on the cardiovascular system. Nat Rev Cardiol. 2018 Mar;15(3): 167–80.

44 Lin LL, Vennarini S, Dimofte A, Ravanelli D, Shillington K, Batra S, et al. Proton beam versus photon beam dose to the heart and left anterior descending artery for left-sided breast cancer. Acta Oncol (Madr). 2015; 54(7): 1032–9.

45 Sardaro A, Petruzzelli MF, D’Errico MP, Grimaldi L, Pili G, Portaluri M. Radiation-induced cardiac damage in early left breast cancer patients: risk factors, biological mechanisms, radiobiology, and dosimetric constraints. Radiother Oncol. 2012 May; 103(2): 133–42.

46 Gori T, Münzel T. Biological effects of low-dose radiation: of harm and hormesis. Eur Heart J. 2012 Feb;33(3): 292–5.

47 Russo GL, Tedesco I, Russo M, Cioppa A, Andreassi MG, Picano E. Cellular adaptive response to chronic radiation exposure in interventional cardiologists. Eur Heart J. 2012; 33(3): 408–14.

48 Borghini A, Gianicolo EAL, Picano E, Andreassi MG. Ionizing radiation and atherosclerosis: current knowledge and future challenges. Atherosclerosis. 2013 Sep;230(1): 40–7.

49 Lee C-L, Blum JM, Kirsch DG. Role of p53 in regulating tissue response to radiation by mechanisms independent of apoptosis. Transl Cancer Res. 2013; 2(5): 412–21.

50 Lee M-O, Song S-H, Jung S, Hur S, Asahara T, Kim H, et al. Effect of ionizing radiation induced damage of endothelial progenitor cells in vascular regeneration. Arterioscler Thromb Vasc Biol. 2012; 32(2): 343–52.

51 Grabham P, Bigelow A, Geard C. DNA damage foci formation and decline in twodimensional monolayers and in three-dimensional human vessel models: Differential effects according to radiation quality. Int J Radiat Biol. 2012; 88(6): 493–500.

52 Girdhani S, Sachs R, Hlatky L. Biological Effects of Proton Radiation: What We Know and Don’t Know. Radiat Res. 2013; 179(3): 257–72.

53 Sasi SP, Song J, Park D, Enderling H, McDonald JT, Gee H, et al. TNF-TNFR2/p75 signaling inhibits early and increases delayed nontargeted effects in bone marrow-derived endothelial progenitor cells. J Biol Chem. 2014; 289(20): 14178–93.

54 Cervelli T, Panetta D, Navarra T, Andreassi MG, Basta G, Galli A, et al. Effects of single and fractionated low-dose irradiation on vascular endothelial cells. Atherosclerosis. 2014; 235(2): 510–8.

55 Bakkenist CJ, Kastan MB. DNA damage activates ATM through intermolecular autophosphorylation and dimer dissociation. Nature. 2003; 421(6922): 499–506.

56 Alan Mitteer R, Wang Y, Shah J, Gordon S, Fager M, Butter P-P, et al. Proton beam radiation induces DNA damage and cell apoptosis in glioma stem cells through reactive oxygen species. Sci Rep. 2015; 5(1): 13961.

57 Lee JH, Koh YA, Cho CK, Lee SJ, Lee YS, Bae S. Identification of a novel ionizing radiation-induced nuclease, AEN, and its functional characterization in apoptosis. Biochem Biophys Res Commun. 2005; 337(1): 39–47.

58 Broustas CG, Xu Y, Harken AD, Chowdhury M, Garty G, Amundson SA. Impact of Neutron Exposure on Global Gene Expression in a Human Peripheral Blood Model. Radiat Res. 2017 ;187(4): 433–40.

59 Tilton SC, Markillie LM, Hays S, Taylor RC, Stenoien DL. Identification of Differential Gene Expression Patterns after Acute Exposure to High and Low Doses of Low-LET Ionizing Radiation in a Reconstituted Human Skin Tissue. Radiat Res. 2016; 186(5): 531–8.

60 Maier P, Hartmann L, Wenz F, Herskind C. Cellular Pathways in Response to Ionizing Radiation and Their Targetability for Tumor Radiosensitization. Int J Mol Sci. 2016 Jan 14; 17(1): 102.

61 Antoccia A, Sgura A, Berardinelli F, Cavinato M, Cherubini R, Gerardi S, et al. Cell cycle perturbations and genotoxic effects in human primary fibroblasts induced by low-energy protons and X/gamma-rays. J Radiat Res. 2009; 50(5): 457–68.

62 Chang PY, Bjornstad K a, Rosen CJ, McNamara MP, Mancini R, Goldstein LE, et al. Effects of iron ions, protons and X rays on human lens cell differentiation. Radiat Res. 2005; 164(4 Pt 2): 531–9.

63 Nam SY, Cho CK, Kim SG. Correlation of increased mortality with the suppression of radiation-inducible microsomal epoxide hydrolase and glutathione S-transferase gene expression by dexamethasone: Effects on vitamin C and E-induced radioprotection. Biochem Pharmacol. 1998; 56(10): 1295–304.

64 Ganapathy V, Smith SB, Prasad PD. SLC19: the folate/thiamine transporter family. Pflugers Arch. 2004 Feb;447(5): 641–6.

65 Zastre JA, Hanberry BS, Sweet RL, McGinnis AC, Venuti KR, Bartlett MG, et al. Up-regulation of vitamin B1 homeostasis genes in breast cancer. J Nutr Biochem. 2013; 24(9): 1616–24.

66 Tiwana GS, Prevo R, Buffa FM, Yu S, Ebner DV, Howarth A, et al. Identification of vitamin B1 metabolism as a tumor-specific radiosensitizing pathway using a high-throughput colony formation screen. Oncotarget. 2015; 6(8): 5978–89.

67 Luo-Owen X, Pecaut MJ, Rizvi A, Gridley DS. Low-dose total-body γ irradiation modulates immune response to acute proton radiation. Radiat Res. 2012; 177(3): 251–64.

68 Kumar P, Miller AI, Polverini PJ. p38 MAPK mediates gamma-irradiation-induced endothelial cell apoptosis, and vascular endothelial growth factor protects endothelial cells through the phosphoinositide 3-kinase-Akt-Bcl-2 pathway. J Biol Chem. 2004; 279(41): 43352–60.

69 Boerma M, Hauer-Jensen M. Preclinical Research into Basic Mechanisms of Radiation-Induced Heart Disease. Cardiol Res Pract. 2010; 2011(858262): 1–8.

70 Streeter PR, Dudley LZ, Fleming WH. Activation of the G-CSF and Flt-3 receptors protects hematopoietic stem cells from lethal irradiation. Exp Hematol. 2003; 31(11): 1119–25.

71 Ossetrova NI, Sandgren DJ, Blakely WF. Protein biomarkers for enhancement of radiation dose and injury assessment in nonhuman primate total-body irradiation model. Radiat Prot Dosimetry. 2014; 159(1–4): 61–76.

72 Soucy KG, Lim HK, Benjo A, Santhanam L, Ryoo S, Shoukas AA, et al. Single exposure gamma-irradiation amplifies xanthine oxidase activity and induces endothelial dysfunction in rat aorta. Radiat Environ Biophys. 2007 Jun;46(2): 179–86.

73 Grabham P, Sharma P. The effects of radiation on angiogenesis. Vasc Cell. 2013 Oct 26;5(1): 19.

74 Girdhani S, Lamont C, Hahnfeldt P, Abdollahi A, Hlatky L. Proton Irradiation Suppresses Angiogenic Genes and Impairs Cell Invasion and Tumor Growth. Radiat Res. 2012; 178(1): 33–45.

